# “The stressed bird in the hand”: Influence of sampling design on the physiological stress response in a free-living songbird

**DOI:** 10.1101/2020.10.05.326694

**Authors:** Nikolaus Huber, Katharina Mahr, Zsófia Tóth, Endre Z. Szarka, Yusuf Ulaş Çınar, Pablo Salmón, Ádám Z. Lendvai

## Abstract

Despite the widely used application of standardized capture-handling protocols to collect blood and assess the physiological stress response, the effect of the actual sampling design (e.g. timing and the number of blood samples) often differs between studies, and the potential implications for the measured physiological endpoints remain understudied. We, therefore, experimentally tested the effects of repeated handling and multiple blood sampling on the stress response in wintering free-living great tits (*Parus major*). We modified a well-established sampling protocol of avian studies by adding either an additional blood sample or a “sham-manipulation” (i.e. handling associated with the blood sampling procedure without venepuncture), to disentangle the effects of handling stress and blood loss. We combined three different stress metrics along the endocrine-immune interface to investigate the acute short-term stress response: total corticosterone levels (CORT), the heterophil/lymphocyte ratio (H:L), and the Leukocyte Coping Capacity (LCC). Our study provided three key results: i) no relationship between CORT-levels, LCC and H:L, confirming that these three parameters represent different physiological endpoints within the stress response; ii) contrasting dynamics in response to stress by the measured parameters and iii) no difference in stress levels 30 minutes after capture due to one additional blood sampling or handling event. By optimising the sampling design, our results provide implications for animal welfare and planning experimental procedures on stress physiology in passerine species.

**Summary Statement:** When testing the short-term stress response in free living passerines, both – the scientist and the bird may be better off with a 15-minute stress protocol.

## Introduction

Sampling blood to assess the individual stress response became a standard procedure in field biology, conservation- and veterinary sciences. In wild, free-ranging animals, however, finding the balance between a reliable representation of physiological conditions by increasing blood sampling units, while minimising handling-induced stress and its effects on individual welfare is a challenging task (Brown and Brown, 2009; de Jong, 2019; Rushen, 1991; Sheriff et al., 2011). In addition, in small-sized vertebrate species, the amount of blood that can be collected is limited and drawing multiple samples over a short time may induce additional physiological changes. Hence, understanding how multiple blood sampling, repeated handling and restrain-time affect individual physiological response patterns is imperative to maintain high standards of animal welfare and might have further implications for the interpretation of physiological target parameters (Johnstone et al., 2012; Owen, 2011; Voss et al., 2010).

Passerine species are a popular model system, and the current standards of good sampling and handling practice in birds have been widely discussed and follow strict rules, *i*.*e*. short handling procedures and blood sampling within 1% of the body-mass (Fair et al., 2010; Owen, 2011). In this context, several studies suggest that taking multiple blood samples from a healthy animal, has minor effects on its condition, behaviour and survival (Ardern et al., 1994; Bonnet et al., 2020; Davis, 2005; Lubjuhn et al., 1998; Stangel, 1986; Voss et al., 2010). Birds, however, perceive the capture and restraint during blood sampling as a life-threatening situation. Given the severity of this acute stress response, the possible effects of the sampling procedure itself (e.g. blood loss) may only marginally affect measurable stress parameters (Bonnet et al., 2020; Davis, 2005; Stangel, 1986; Zylberberg, 2015). Whereas it is well recognised that stressors (i.e. unpredictable, uncontrollable adverse changes in the environment) experienced before capture, may have a significant impact on the measurements of individual physiological stress responses, only a few studies have shown that even after the onset of handling-stress, additional disturbance can alter the physiological response patterns (Cockrem, 2013; Gratto-Trevor et al., 1991; Pakkala et al., 2013; Wingfield et al., 1995). For example, Canoine et al. (2002), reported higher stress-associated hormone levels (corticosterone; CORT) in birds exposed to a predator in addition to the blood sampling procedure and thereby demonstrated a surprisingly high context-specific plasticity in the stress response. On the contrary, studies which applied different stress indicators, such as the change in total white blood cells (WBC), show that additional handling and blood sampling only has negligible effects (Bonnet et al., 2020; Davis, 2005). A possible explanation for this discrepancy may be that the different physiological stress systems and their measurable endpoints vary in their sensitivity to stressors and the timescale each parameter integrates stress (Gormally and Romero, 2020). These differences potentially impede the interpretation of data, aiming to reveal the effects of multiple blood sampling and handling on the individual stress response and the potential direct effects of sampling (e.g. blood loss) (Davis, 2005; Sheldon et al., 2008; Voss et al., 2010).

We, therefore, tested different blood sampling regimes and their impact on the outcome of short-term stress in a free-living passerine species, the great tit (*Parus major*). We used the well-established “capture-handling stress” protocol (Wingfield et al., 1982), as the underlying framework and included three stress metrics representing different physiological systems with varying response latencies towards handling-induced stress (Davis and Maney, 2018; Gormally and Romero, 2020; Johnstone et al., 2012). The basic protocol is designed to measure the activation and reactivity of the Hypothalamic-Pituitary-Adrenal- (HPA-) axis and the concomitant changes in circulating CORT levels in response to a defined stressor (i.e. capture and handling) by collecting a series of blood samples. The most commonly applied form of the protocol (we refer to it as “standard protocol”) is a reduced version of the pioneering study by Wingfield et al., 1982 and includes the collection of a sample immediately after capture (< 3 min), which is regarded to represent CORT levels near the baseline, and 30 min thereafter, considered as the CORT-peak response (Romero and Reed, 2005; Romero and Wingfield, 2016).

CORT levels increase within minutes after the onset of a stressor and act on several physiological pathways in a parallel manner to adjust bio-regulatory mechanisms and enhance immediate survival. These effects involve the support of essential physiological functions (e.g. cardiac-, respiratory and brain-activity), energy mobilisation and re-establishment of homeostasis after a stressful event (Sapolsky et al., 2000). Stress responses also affect the distribution and function of innate immune cells (Dhabhar, 2002; Verburg-van Kemenade et al., 2017). Whereas lymphocytes migrate from the circulation into the peripheral tissues, polymorphonuclear granulocytes (PMNLs, i.e. neutrophil granulocytes in mammals and heterophil granulocytes in birds) diffuse from the periphery into the bloodstream. Therefore the ratio between heterophil granulocytes and lymphocytes (H:L ratio) became a reliable and frequently used measure for stress in birds and was also incorporated into our study-design (Davis and Maney, 2018; Davis et al., 2008). Further, we included a rather recently established parameter, the leukocyte coping capacity (LCC), to measure an additional aspect of the immunological stress response (McLaren et al., 2003). In the course of a stress response, the glucocorticoid-, α- and β-adrenoceptors of heterophil granulocytes are activated and an “oxidative burst”, is triggered, leading to the release of superoxide free radicals and the production of reactive oxygen species (ROS) (Frolov et al., 2006; Ronchetti et al., 2018). This reaction can be simulated under experimental conditions and measured in real-time via chemiluminescence from whole blood (Huber et al., 2020; Huber et al., 2019).

In our experiment we applied modified versions of the “standard protocol” and assigned individual great tits to three sampling regimes: (1) *Standard*, i.e. blood samples at baseline (<3 min) and 30 min post-capture; (2) *3-samples*, where we included an additional blood sample 15 min post-capture; (3) *Sham*, i.e. handling without venepuncture at 15 min post-capture, in order to separate the potential effects of blood loss from handling. If stress intensity affects the measured physiological parameters, we predict higher CORT levels in individuals undergoing higher handling frequency (groups 2 and 3), in comparison to individuals assigned to the *Standard* protocol. Considering that heterophil numbers increase in the bloodstream after a stressor, we also expect that the H:L ratio will be elevated in the groups with additional blood sampling and/or handling, i.e. *3-samples* and *Sham*. However, if blood loss and hence the decrease of blood cells affect these measures, the H:L ratio is expected to decrease in birds in the *3-samples* group. Changes in the H:L ratio are considered to be slow in comparison to the other parameters, but 30 min after the onset of the stressor will be sufficient to reveal differences in comparison to baseline levels (Davis, 2005). Regarding the LCC dynamics, we predicted that birds undergoing the *Standard* would show an increase in LCC levels with the time of the procedure (i.e. a partial recovery from capture and the first handling in the cloth bag). In contrast, individuals from the *3-samples* and *Sham* protocols will show a decreasing dynamic in LCC levels over the timespan of the experiment, which is expected to be even more pronounced in the *3-samples* group.

## Methods

We caught 58 great tits (*Parus major*) on a winter-feeding site in the Botanical Garden of the University of Debrecen (47°33’24.6”N, 21°37’16.2”E). The artificial feeder was established one week prior to the start of the experiment and regularly maintained. To reach a sufficient sample size, we provided an additional stimulus: we placed a speaker under the feeder, which broadcasted the calls of a mixed-species flock (in addition to great tit, the playback included blue tit, *Cyanistes caeruleus*, long-tailed tit *Aegithalus caudatus* and coal tit *Periparus ater*). This playback type is considered to attract birds without causing a territorial/stress response.

Birds were captured using mist nets between December 2018 and February 2019. All samples were collected between 09:00-12:00. We collected a blood sample immediately after the capture by puncturing the brachial vein using a sterile 26G needle. The initial blood sample (∼50 μl) was drawn into heparinised micro capillary-tubes immediately after a bird hit the net, (average time in s: 155 ± 51 (SD) s, range: 61-291 s). Immediately after the sampling, the birds received a numbered aluminium ring and were transferred into a cotton bag. Individuals were randomly assigned to the three treatment groups after the first sample was drawn. In the first group (*Standard*), after initial sampling, the birds remained in a cloth bag until the second sample was drawn. In the second group (*3-samples*), we collected blood samples, 15 min and 30 min after the capture. In the third experimental group (*Sham*), we took blood samples at <3 (hereafter baseline) and 30 minutes as in the *Standard* group, but in order to separate the effects of handling from blood loss, 15 min after the capture we performed the entire handling procedure (removing the bird from the bag, opening and preparing the wing for blood sampling) without the actual venepuncture.

Immediately after each sampling event, a small drop of blood was used to prepare the blood smears using the two-slide wedge technique. In addition, we aliquoted 20 µl whole blood for the LCC analyses, and the remaining sample was centrifuged for 5 minutes to separate plasma from red blood cells. We removed the plasma with a Hamilton syringe and stored the samples in a –20°C freezer until assayed for corticosterone.

### Leukocyte coping capacity

Immediately after blood collection, 20 µl of heparinised whole blood was transferred into a silicon anti-reflective tube (Lumivial, EG & G Berthold, Germany), containing 180 µl of 10^−4^ mol l^−1^ lucigenin (bis-N-methylacridinium nitrate; Sigma Aldrich, Vienna, Austria) dissolved in dimethyl sulfoxide (DMSO; VWR International, Stockholm, Sweden) and diluted in Phosphate Buffered Saline (PBS, pH 7.4). Lucigenin produces chemiluminescence when combined with an oxidising agent (i.e. superoxide anion) and was used to quantify the production of extracellular reactive oxygen species (ROS) production of heterophil granulocytes in real-time (Gyllenhammar, 1987; Li et al., 1998). After that, the sample was mixed gently and aliquoted into two anti-reflective tubes. In one tube, 10 µl of PBS (pH 7.4) was added in order to measure unstimulated blood chemiluminescence levels, providing information on individual baseline levels of superoxide anion and acts as a control. In the second tube, we added 10 µl of 10^−5^ mol l^−1^ phorbol-myristate-acetate (PMA; Sigma Aldrich, Vienna, Austria) to assess full blood chemiluminescence in response to this secondary (the first natural challenge occurs in vivo) chemical challenge (Shelton-Rayner et al., 2012). Blood chemiluminescence for each tube was measured for 30 seconds every 10 minutes over 80 minutes by using a portable high sensitivity chemiluminometer (Junior LB 9509, EG & G Berthold, Germany) and is expressed in relative light units (RLU; arbitrary scale reflecting photon count divided by 10). All measurements were carried out in the laboratory with temperatures between 20°C and 25°C. The tubes were kept at 40°C in a glass beaker with metal beads placed in a water bath (40°C) to simulate *in vivo* temperature conditions slightly below the average active body temperature and were gently swivelled from time to time to avoid pelleting of the blood cells (Prinzinger et al., 1991). Samples were not centrifuged as the texture and adhesiveness of the cell microenvironment is essential for the in vivo determination of cell reactivity (Théry et al., 2007). In order to correct for background noise, we subtracted the values of the control sample from that of the challenged sample measured at the same time point. From the resulting LCC response curve reflecting the PMA induced production of PMNL ROS in real-time, we extracted the LCC peak as a variable. To account for individual differences in the number of heterophils and a potential mass effect on LCC, we used the residuals of a general additive model between the LCC peak and the number of heterophils (Fig. S1), hereafter ‘corrected LCC’ as our response variable in the analyses (see *Statistical analyses*).

### Leukocyte cell count

Blood smears were air-dried, fixed with ethanol and dyed with Wright-Giemsa Quick stain following previous protocols established for songbirds (e.g. Cīrule et al., 2012). Briefly, smears were examined at 1,000× magnification, and a minimum of 50 leukocytes was counted per slide while keeping track of the number of view fields and the number of erythrocytes. The number of each leukocyte type, i.e. lymphocytes, heterophils, eosinophils, monocytes and basophils, was expressed per 10,000 erythrocytes. In addition, the total number of white blood cells and the heterophil to lymphocyte ratio (H:L) was calculated (Pap et al., 2015). All the cell counts were performed by the same observer (E.Z. Szarka), and a random subset of smears (n = 15) was analysed in duplicates showing a moderately to high repeatability between counts for the main cellular types, lymphocytes (R = 0.722; 95% CI [0.349, 0.891]; p< 0.001), heterophils (R = 0.492; 95% CI [0.029, 0.817]; p = 0.028) and the H:L ratio (R= 0.84; 95% CI[0.592, 0.942]; p< 0.001).

### Hormone assays

We quantified the total corticosterone, using direct radioimmunoassay (Lendvai et al., 2011). Before the assays, we extracted the corticosterone from the plasma samples, using diethyl-ether. Then, the extracts were reconstituted in PBS. After overnight incubation at 4 °C, we added ∼10K dpm of 3H-Cort (Catalogue number: NET399250UC, lot number: B00025; Perkin Elmer, Waltham, MA, USA), antiserum (MP Biomedicals 07-120016, lot number: 3R3-PB-20E2) and PBS. After another incubation overnight at 4°C, the dextran-coated charcoal was added to separate corticosterone bound to antibodies. The radioactivity of the bound fraction was counted in a liquid scintillation counter (QuantaSmart). We processed all samples in one assay (intra-assay CV = 4.17%).

### Statistical analysis

The physiological short-term stress response was analysed from 57 individuals. All statistical analyses were conducted using R version 3.6.2 (R Core Team, 2019). Physiological variables were analysed in linear mixed-effects models using treatment group and sampling time points (baseline, 15 and 30 min) as fixed effects (factors) and individual identity as a random intercept. The models also included the group × sampling point two-way interaction, as this was part of the experimental design. Degrees of freedom were determined using Satterthwaite’s approximation and significance tests were obtained as implemented in the ‘lmerTest’ R package (Kuznetsova et al., 2017). Assumptions of the models were assessed by visual inspection of the residuals. H:L was arc-sine square-root transformed. CORT and LCC values were not transformed. To test the relationship between the variables recorded to represent different aspects of the individual short-term stress response, we conducted principal component analysis (PCA) on H:L, LCC, and CORT levels.

## Results

### Relationship between the response variables

At the baseline level (i.e. < 3min), principal component analyses showed that CORT, LCC and H:L were not strongly related. The PCA had three components, with Eigenvalues PC1 = 1.12, PC2 = 0.97, PC3 = 0.89. The variance explained by each component was similar (PC1 = 41.6%, PC2 = 31.5%, PC3 = 26.8%; Fig 1a). While all variables loaded positively in PC1, LCC and H:L seemed to represented an independent axis to CORT in the second component, although still three variables independent from each other (Fig. 1a, PC2). Short-term stress (i.e. capture and subsequent handling and constraint for 30 min) did not modify the relationship between the three response variables and they remained largely independent from one another (Fig. 1b). Eigenvalues of the PCA were: PC1 = 1.11, PC2 = 1.04, PC3 = 0.82. The variance explained by each component was similar to the baseline case: (PC1 = 41.1%, PC2 = 36.2%, PC3 = 22.7%).

**Fig. 1.**
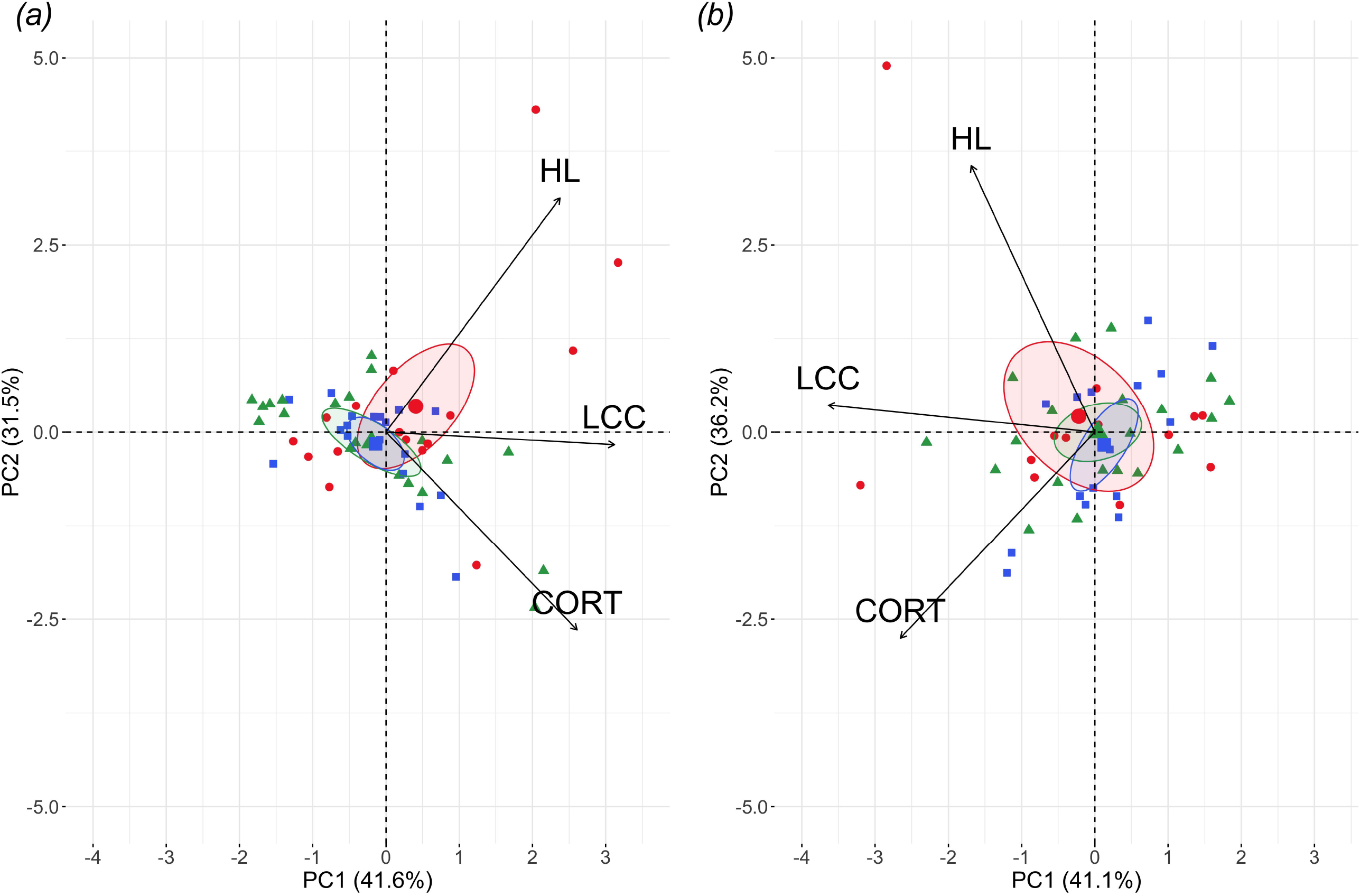
Principal component analysis of the three physiological parameters: total CORT, LCC (corrected by heterophils number) and H:L-ratio (HL) measured (a) at baseline (< 3 min) or (b) after 30 minutes. In the course of the stress response, the three response variables where re-organised but remained largely independent from one another, showing that these stress indicators represent different physiological aspects within the stress response. Likewise, the variation explained by each component was similar to the baseline case. Small points represent each individual’s sampling point while the bigger symbols indicate the bivariate median response with the 95% CI ellipse. *Standard* (red circles), *Sham* (green triangles) and *3-samples* (blue squares).

### Effects of the blood sampling regime on the three stress response variables

Baseline CORT levels did not differ between the three sampling regimes (F_2,63_ = 0.79, p = 0.456). Capture and handling stress induced a significant increase of corticosterone (F_2,69_ = 54.61, p < 0.001), both at 15 minutes (t = 4.34, p < 0.001) and at 30 minutes (t = 6.24, p < 0.001) post capture. However, at 30 minutes there was no difference between the three experimental groups (F_2,69_ = 0.44, p = 0.645): corticosterone levels in the *Standard* group did not differ from either the *Sham* (t = –0.79, p = 0.432) or the *3-samples* (t = –0.86, p = 0.391, Fig. 2a, Fig. 3a).

**Fig. 2.**
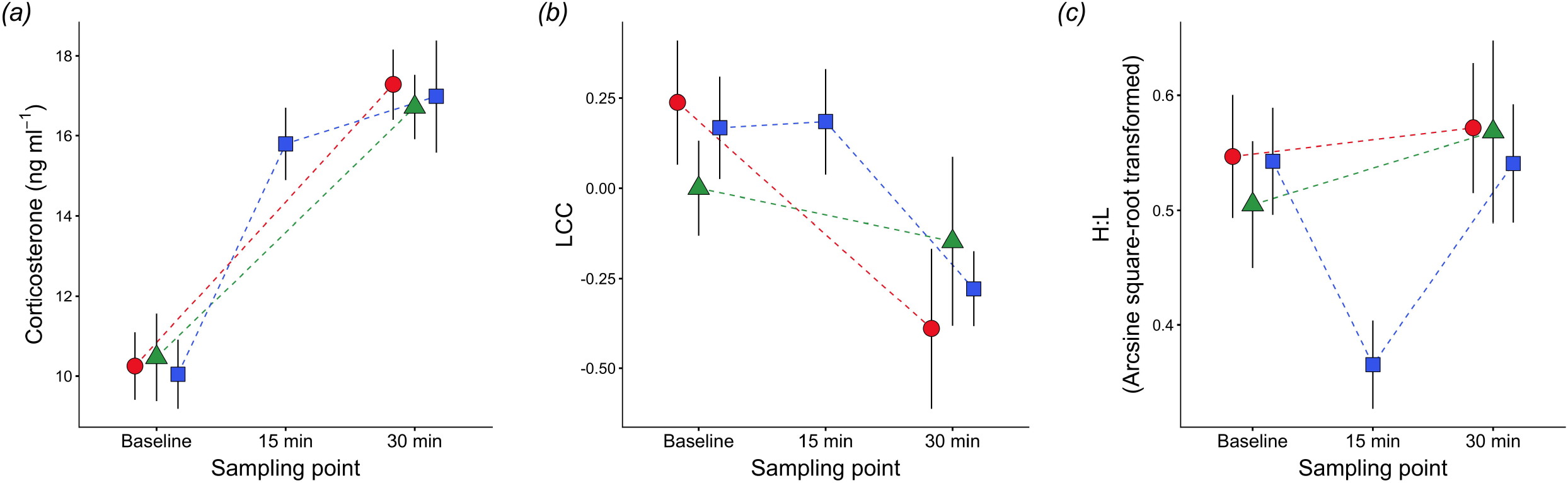
Physiological levels of wild great tits expose to the three different handling regimes, *Standard* (red circles), *Sham* (green triangles) and *3-samples* (blue squares). (a) Total corticosterone levels (CORT); (b) Leucocyte Coping Capacity (LCC, corrected by heterophils number) and (c) H:L (arcsine square-root transformed). Symbols represent means ± standard error of the means at baseline (< 3 min), 15 min and 30 min after capture.

**Fig. 3.**
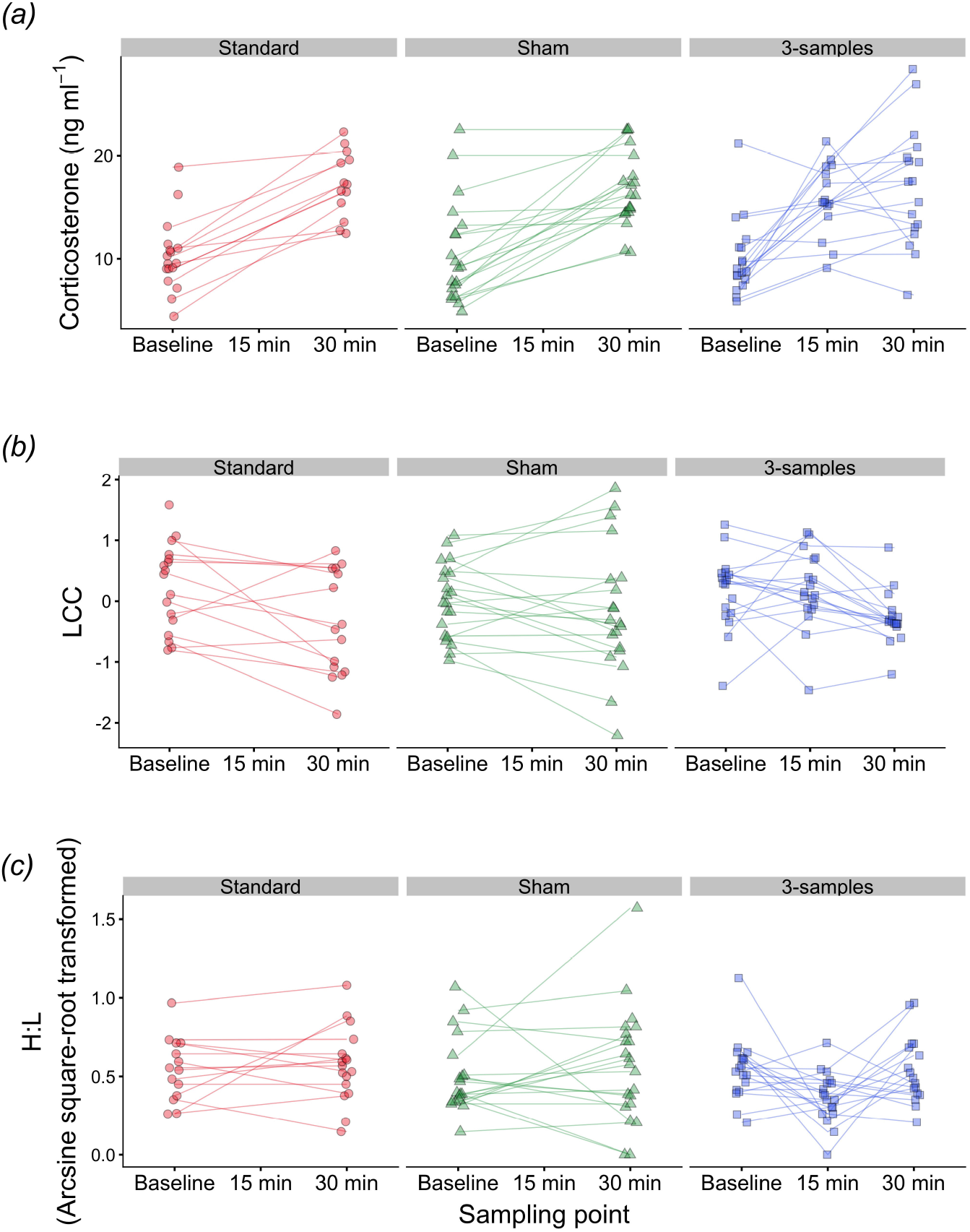
Individual physiological responses of wild great tits expose to the three different handling regimes, *Standard* (red circles), *Sham* (green triangles) and *3-samples* (blue squares). (a) Total corticosterone levels (CORT); (b) Leucocyte Coping Capacity levels (LCC, corrected by heterophils number) and (c) H:L (arcsine square-root transformed). Symbols represent each individual and lines connect the measurements of the same individual at each time point: baseline (< 3 min), 15 min and 30 min after capture.

LCC levels at capture did not differ between the three groups (F_2,56_ = 0.10, p = 0.902). In contrast to the CORT response, capture and handling induced a drop in LCC values (F_2,66_ = 5.96, p = 0.004), but only after 30 minutes (t = –3.04, p = 0.003), not at 15 minutes (t = 0.50, p = 0.620) post capture. The experimental groups did not differ in their LCC levels at 30 minutes (F_2,67_ = 0.69, p = 0.505, Fig. 2b, Fig. 3b).

Similarly to the other two parameters, we observed no difference in the baseline H:L (F_2,55_ = 0.05, p = 0.956) between the three protocols. In the *3-samples* group, handling induced a significant drop in H:L (F_2,59_ = 3.83, p = 0.027) after 15 minutes (t = –2.39, p = 0.02), but H:L values returned to the baseline levels after 30 minutes, and thus did not differ from the *Standard* group (t = –0.29, p = 0.772). Also, H:L in the *Sham* group did not differ from the *Standard* group after 30 minutes (t = 0.27, p = 0.791, Fig. 2c, Fig 3c).

## Discussion

Our study provides three key results. First, in line with previous findings, we reveal only a weak relationship between CORT, H:L and LCC (Fig. 1), confirming that these three parameters represent different physiological aspects within an acute, short-term stress response (Davis and Maney, 2018; Gormally et al., 2019; Huber et al., 2017). Second, we found that these three physiological parameters showed different kinetic dynamics over the time course of the short-term stress response, reflecting different response latencies (Fig. 2) (Gormally and Romero, 2020). CORT levels increased markedly in most individuals, but the timing of the peak response underlies substantial between-individuals variation (Fig. 3a). On the contrary, the LCC response decreased steadily, with no noticeable differences regarding the timing of the blood sample (Fig. 2b, Fig. 3b). However, our results also revealed a sudden drop of the H:L ratio 15 min post-capture with a subsequent recovery to baseline levels 30 min post-capture (Fig. 2c, Fig. 3c). Finally, we showed that the different sampling procedures applied in this study did not lead to significant differences in the CORT, H:L ratio and LCC response 30 min after capture.

We predicted a stronger HPA-axis activation with higher CORT levels in birds that experienced additional handling stress and underwent three sampling procedures. This effect, however, did not become apparent in our study which is supported by findings in other avian and non-avian species (Bonnet et al., 2020; Davis, 2005; Stangel, 1986). Bonnet and colleagues, for example, recently reported, that blood sampling *per se* did not affect CORT levels in dice snakes (*Natrix tessellata*) (Bonnet et al., 2020). Whereas individual condition (i.e. health or nutritional status) and previously experienced stress events can change the stress responsiveness and the overall HPA-axis response, additional handling after the onset of the initial stress stimulus may have only little effects on CORT levels. Circulating CORT levels increased in almost all birds between baseline and 15 min, with some individuals showing a weak further increase or even a decrease after the second bleeding, possibly as the result of the effective negative feedback of CORT within the HPA axis. These results are corroborated by previous work on songbirds (Romero and Remage-Healey, 2000), including recent studies on migrating garden warblers (*Sylvia borin*) and great tits, suggesting that in some species measuring CORT at 15 min (or 10 min) post-capture may be sufficient (and arguably preferable) to study the magnitude of HPA axis activation towards standardised acute, short-term stress (Cockrem, 2013; Huber et al., 2020).

We observed a similar pattern when exploring the effects of the experimental protocol on the shift in leukocyte profiles (Romero and Wingfield, 2016; Skwarska, 2019). The activation of the HPA axis and the consecutive increase of circulating CORT is leading to a redistribution of innate immune cells into specific target tissues and thereby to a reduction of circulating lymphocytes while simultaneously increasing the number of circulating heterophils (Cīrule et al., 2012; Lentfer et al., 2015; Maxwell, 1993; Sturkie, 2015). This relationship was previously demonstrated in CORT supplementation experiments, but also capture and restraint stress was shown to be sufficient to effect similar responses as is corroborated by a growing number of studies on vertebrates (Cīrule et al., 2012; Davis et al., 2008; Lentfer et al., 2015). The H:L ratio, however, is considered to mainly reflect long-term, rather than an immediate, short-term physiological stress response (Gormally and Romero, 2020; Maxwell, 1993; O’Dell et al., 2014) and there is a strong indication that an increase after capture becomes significant only 30 to 60 min post-capture and in a species-specific manner (Davis, 2005; Davis and Maney, 2018). Remarkably, our data show an unexpectedly fast decrease in the H:L ratio at 15 min after capture, followed by rapid recovery to baseline levels 30 min post-capture. One possible explanation is the acute “*fight-and-flight*” response and the concomitant increase in noradrenaline, causing the observed transient lymphocytosis shortly after capture (Eriksson and Hedfors, 1977; Ince et al., 2019). Furthermore, severe stress may cause the exhaustion of mature cell pools and the release of immature cells from the bone marrow, which can result in transient heteropenia (Lentfer et al., 2015; Maxwell, 1993; Romero and Wingfield, 2016). Hence, the rapid drop and the subsequent recovery of the H:L ratio measured in our samples may be due to short-term physiological compensatory mechanisms of the organism in response to severe handling stress and possibly also the sudden loss of blood.

Interestingly, LCC levels did not vary significantly between sampling regimes and decreased in all groups equally in the course of the experiment. LCC has been suggested as a reliable and sensitive tool to measure stress in mammals and birds and has been successfully applied to assess psychological stress in humans (Mian and MacDonald, 2010; Shelton-Rayner et al., 2012). In respect to the existing literature, a decrease in the LCC response, or a lack of recovery respectively, is an indicator for high stress levels and a reduced capacity to cope with and/or recover from a stress event (Huber et al., 2019; McLaren et al., 2003). For instance, in captive house sparrows (*Passer domesticus*), leaving the birds undisturbed for 30 min in a cotton bag was sufficient to allow individuals to recover from capture stress in winter, reflecting in an increase of the measured LCC response (Huber et al., 2017). Based on these findings, we expected a more definite decline in the LCC response in birds that underwent multiple sampling and handling and have hence experienced a disrupted recovery period. Our results, however, show that LCC decreased in all three experimental groups and neither the additional handling nor the third blood sample resulted in a different LCC response between the experimental groups 30 min post-capture. However, the captive house sparrows in Huber et al. (2017) had *ad libitum* access to food and were habituated to human presence. These conditions may have positively contributed to the recovery of the LCC response and a faster re-establishment of homeostasis after capture and handling (Gormally et al., 2019). Wintering great tits face harsh weather conditions and limited access to food, possibly reducing the capacity to recover from the capture/handling stress. It should also be noted that LCC levels assessed from the additional blood sample after 15 min did not significantly differ from the first sample and only decreased thereafter to a similar level as observed in the other two groups. Our results also suggest that the actual blood sampling (loss) had no effect since LCC levels in the *3-samples* group at 30 min are not different from the *Sham* or the *Standard* group. However, we already controlled for a potential mass effect of the changing white blood cell composition on LCC by correcting this measurement for the individual number of heterophils (corrected LCC). The number of studies on LCC in passerines is limited (Huber et al., 2020; Huber et al., 2017), and the observed patterns might be species-specific and underlie seasonal variation. Furthermore, Gormally et al. (2019) have demonstrated that HPA axis regulation, immune function, antioxidant levels, DNA damage and stress-related alterations in behaviour change on distinct timescales in response to repeated stressors or differences in recovery time between the stress events (Gormally and Romero, 2020). We would, therefore, like to emphasise that we only recorded data over a short time frame and encourage follow-up studies to measure the recovery from capture and handling induced stress over a more extended period. Moreover, additional measures for stress, representing different physiological aspects within the stress response such as oxidative stress, should be applied and tested in future studies (Costantini, 2008; Stier et al., 2019).

The sharp drop of the H:L ratio and more importantly, the fast recovery to baseline levels 30 min post-capture might provide an interesting new perspective for future research questions. Our results also revealed a decrease of the LCC response only after 15 to 30 min post-capture, and we speculate that a resembling pattern may have occurred in the *Standard* and *Sham* groups. The observed response in the *3-samples* group indicates that birds did not recover from the initial capture stress after 15 min, and their condition in terms of LCC deteriorated until 30 min after capture. Even though it is commonly assumed that keeping the birds in a dark cotton bag reduces stress and might even lead to a recovery, it should be considered that restraining a wild animal over a specific time-limit can result in severe stress and pathology (Breed et al., 2020). Thus, shortening the period of human presence and restrain, and reducing handling frequency, may be the only sufficient method to minimise additional stress imposed on the animals. This is of particular importance during specific life-history stages, such for example as reproductive period, moulting or migration, where animals are more vulnerable towards stressors. Besides, the prolonged removal of an individual from its natural habitat might have other downstream consequences for an individual’s fitness and/or performance such as lower foraging success, decreased food provisioning for nestlings or even nest desertion. We acknowledge that stress responses are likely to be species-specific, and not all experimental procedures allow fast processing of the animals and require extended periods of capture and restrain. However, we advocate the investigation of shorter sampling regimes and its preferable utilisation over the standard 30 min when it seems suitable, for both biological and ethical reasons.

## Author contributions

*Conceptualization*: N.H, K.M., A.Z.L.; *Methodology:* K.M., Z.T., E.S., Y.U.C., N.H.; *Formal analysis*: A.Z.L., P.S.; *Writing - original draft*: K.M., N.H.; *Review & editing*: P.S., Z.T., E.S., Y.U.C., A.Z.L.

## Acknowledgements

We would like to thank A. Fülöp, G. Rédai and L. Őri for their help during data collection and maintaining the feeders as well R. Hengsberger, for her help with literature search and copy-editing. Special thanks to DVM Erika Fuhrmann for her input and discussions on dynamics in blood cell compositions.

## Ethics

We followed all applicable international, national, and institutional guidelines for the use of animals. All procedures performed in studies involving animals were in accordance with the ethical standards of the institution and approved by the institutional animal care and use committee.

## Data availability

The raw data supporting the conclusions of this manuscript will be made available by the authors, without undue reservation, to any qualified researcher upon request.

---

**Fig. S1**. General additive model between individual’s Leucocyte Coping Capacity (LCC) and number of heterophils. Both variables were log10-transformed. Coloured points indicate the handling regime: *Standard* (red circles), *Sham* (green triangles) and *3-samples* (blue squares). The black line represents the significant relationship between both variables (F_1, 122_= 25.07, p< 0.001).

